# Basins of attraction of microbiome structure and soil ecosystem functions

**DOI:** 10.1101/2022.08.23.505048

**Authors:** Hiroaki Fujita, Shigenobu Yoshida, Kenta Suzuki, Hirokazu Toju

**Author notes:** **Competing interest:** HT is the founder and director of Sunlit Seedlings Ltd. The other authors declare that the research was conducted in the absence of any commercial or financial relationships that could be construed as a potential conflict of interest.

## Abstract

Theory predicts that biological communities can have multiple basins of attraction in terms of their species/taxonomic compositions. The presence of such basins of community structure has been examined in classic empirical studies on forest–savanna transitions and those on eutrophication in freshwater lakes. Nonetheless, it remains a major challenge to extend the investigations of multistability to species-rich microbial communities. By targeting soil microbiomes, we infer the stability landscapes of community structure based on the concepts of statistical physics. Our analysis on the compiled dataset involving 11 archaeal, 332 bacterial, and 240 fungal families detected from > 1,500 agroecosystem soil samples suggested that both prokaryotic and fungal community compositions could be classified into several basins of attraction. We also found that the basins differed greatly in their associations with crop disease prevalence in agroecosystems. A further analysis highlighted microbial taxa potentially playing key roles in transitions between basins with different ecosystem-scale functions. The statistical framework commonly applicable to diverse microbial and non-microbial communities will reorganize our understanding of relationship among community structure, stability, and functions.

## Introduction

The idea that the state space of biological community structure can comprise multiple basins of attraction have inspired both empirical and theoretical ecologists since the late 1960s (Beisner et al., 2003; May, 1977; Scheffer et al., 1993). The concept of multistability has been examined in aquatic and terrestrial ecosystems (Scheffer and Carpenter, 2003; Schröder et al., 2005; Suding and Hobbs, 2009). Community structure in shallow lakes, for example, is known to show two discrete states depending on nutrient (phosphorus) concentrations as represented by the bistability of charophyte densities (Ibelings et al., 2007; Scheffer et al., 1993; Smith and Schindler, 2009).

Likewise, worldwide inventories of tree cover have shown that forest and savanna vegetation types possibly represent different basins of attraction (Hirota et al., 2011; Staver et al., 2011b, 2011a). Those studies have further shown that ecosystem functions (e.g., fishery, agricultural, and forestry production) can differ greatly between such basins of biological community structure (Gunderson, 2000; Scheffer et al., 2001, 1993; Scheffer and Carpenter, 2003). Consequently, understanding how structure, stability, and biological functions are organized in real communities and ecosystems has been one of the major goals in ecology.

While classic studies targeting freshwater and terrestrial biomes have explored basins of attraction based on simple characterization of community states (e.g., tree cover percentages), recent technical advances in microbial community (microbiome) research have come to provide opportunities for deepening our knowledge of biological community stability (Amor et al., 2020; Costea et al., 2017; Faust et al., 2015; Shaw et al., 2019; Toju et al., 2018; Zaneveld et al., 2017). Based on amplicon and shotgun sequencing technologies, large datasets of microbial species/taxonomic compositions have been made available, providing a basis for exploring reproducible states in microbiome community structure (Amor et al., 2020; H Fujita et al., 2023; Hayashi et al., 2024). Such high-throughput DNA sequencing studies in medicine, for example, have shown that human individuals can be classified into three or four semi-discrete clusters in terms of their intestinal microbiome compositions (Arumugam et al., 2011; Wu et al., 2011) [see also (Jeffery et al., 2012; Knights et al., 2014)]. Intriguingly, these alternative gut microbiomes (“enterotypes”) differ in their associations with human disease such as type II diabetes and Crohn’s disease (Costea et al., 2017). In addition to those studies on animal-associated microbiomes (Arumugam et al., 2011; Moeller et al., 2012; Yajima et al., 2023), studies on plant-associated microbiomes have started to reorganize our recognition of how multistability of phyllosphere/rhizosphere microbiome structure is associated with ecosystem-scale processes and functions (Toju et al., 2018, 2016). Because hundreds or thousands of replicate community samples are available in such microbiome studies, it is now possible to discuss potential relationship between community structure and ecosystem functions based on statistical signs of the presence of multiple basins (and background attractors).

In theoretical ecology, stability of community states (taxonomic or species compositions) is often discussed in the framework of stability landscapes (Beisner et al., 2003; Hastings et al., 2018; Lewontin, 1969; Scheffer and Carpenter, 2003; Suzuki et al., 2021). On the landscape representing stability/instability of community structure, basins of attraction are split by “tipping points” representing unstable equilibria (Beisner et al., 2003; Scheffer et al., 2001; Scheffer and Carpenter, 2003; Suzuki et al., 2021) (Figure 1). As these basins differ in the biological functions of constituent communities, stable and highly functional community states can be explored within the stability landscapes. With the application of a recently proposed mathematical approach developed based on statistical physics (Becker and Karplus, 1997; Watanabe et al., 2014), it is now possible to infer “energy landscapes”, which represent structure of stability landscapes, from empirical datasets of ecological communities (Dakos and Kéfi, 2022; Sánchez-Pinillos et al., 2024; Suzuki et al., 2021). The statistical framework allows us to explore the probabilities of community compositions within the “assembly graphs” (Coyte et al., 2021; Serván and Allesina, 2021) representing paths of possible community assembly (H Fujita et al., 2023; Suzuki et al., 2021) (Figure 1). Although hundreds or thousands of community compositional data points are required to apply the statistical approach (H Fujita et al., 2023; Suzuki et al., 2021), such energy landscape analyses will allow us to define key features of stable and highly functional microbiome states out of numerous possible combinations of microbial species or taxa. Despite the potential for systematically profiling the relationship among community structure, stability, and functions based on massive community datasets, the energy landscape analysis has been applied only to a few microbial community datasets (H Fujita et al., 2023; Suzuki et al., 2021).

**Figure 1.**
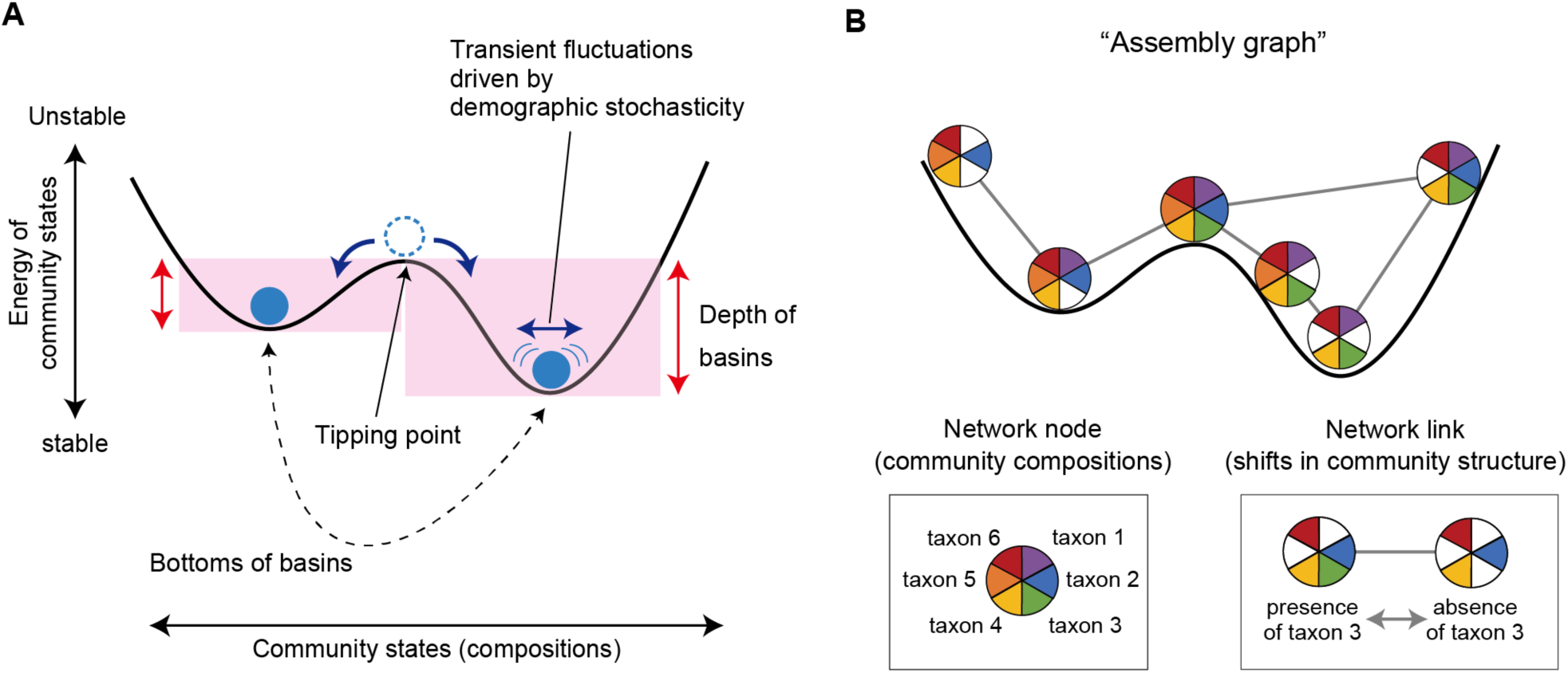
Schema of multistability of ecological communities. (**A**) Basins of attraction and tipping points. The structure of “stability landscapes” showing relationship between community states (species or taxonomic compositions) and their stability is inferred based on the energy landscape analysis. The “energy” of each community state is calculated with maximum entropy models as detailed in Methods. Lower energy represents a more stable community state on a stability landscape. Transient fluctuations around the bottoms of basins (i.e., point attractors) are assumed as probabilistic phenomena in the statistical approach. (**B**) Assembly graph. To explore numerous possible states of real ecological communities, input data are binarized in the energy landscape analysis. Potential transitions between community states are then considered within “assembly graphs”, in which paths between different species/taxonomic compositions are treated as network links. Thus, by the assembly-graph approach, the energy landscape analysis provides a general framework for inferring the structure of stability landscapes in empirical studies of complex microbiome datasets.

We here apply the emerging statistical framework to soil microbiomes, which often show highest levels of structural diversity in nature. We compile a cropland soil microbiome dataset consisting of > 1,500 sampling positions across the Japan Archipelago (Fujita et al., 2024). With the massive dataset, we infer the compositional stability of prokaryotic and fungal communities based on maximum entropy models of the energy landscape analysis (Suzuki et al., 2021). We then examine whether the basins of attraction of soil microbiomes can differ in ecosystem-scale functions by focusing on potential relationship between soil microbial compositions and the prevalence of crop plant disease. We also explore key microbial taxa whose abundance critically divide the basins representing favorable and unfavorable ecosystem functions. The results of the energy landscape analysis are further used to infer tipping points splitting the inferred basins. Overall, this study illustrates how we can integrate the information of community structure, stability, and functions based on a statistical platform commonly applicable to diverse microbial and non-microbial communities.

## Methods

### Dataset compilation

We compiled a publicly available dataset of cropland soil microbiomes (DDBJ accession: DRA015491; Figure 2) with its metadata of the samples (Fujita et al., 2024). In the previous study reporting the data (Fujita et al., 2024), 2,903 bulk soil samples collected from the field of 19 crop plant species (apple, broccoli, cabbage, celery, Chinese cabbage, eggplant, ginger, komatsuna, lettuce, onion, potato, radish, rice, satsuma mandarin, soybean, spinach, strawberry, sweet corn, tomato) across the Japan Archipelago from January 23, 2006 to July 28, 2014 (latitudes of the sampling positions: 26.1–42.8 °N) were subjected to the amplicon sequencing analysis of the prokaryotic 16S rRNA region and the fungal internal transcribed spacer 1 (ITS1) region (Fujita et al., 2024). The information of dry soil pH, electrical conductivity, carbon/nitrogen (C/N) ratio, and available phosphorous concentration was available for 2,830, 2,610, 2,346, and 2,249 samples, respectively. Likewise, the information of crop plant disease [the percentage of diseased plants or disease severity index (Chiang et al., 2017)] was available for 1,471 samples (Fujita et al., 2024). The plant pathogens surveyed were *Colletotrichum gloeosporioides* on the strawberry, *Fusarium oxysporum* on the celery, the lettuce, the strawberry, and the tomato, *Phytophthora sojae* on the soybean, *Plasmodiophora brassicae* on Cruciferae plants, *Pyrenochaeta lycopersici* on the tomato, *Pythium myriotylum* on the ginger, *Ralstonia solanacearum* on the eggplant and the tomato, and *Verticillium* spp. on Chinese cabbage (Fujita et al., 2024). After a series of quality filtering, prokaryotic and fungal community data were available for 2,318 and 2,186 samples, respectively. In total, 579 archaeal amplicon sequence variants (ASVs) representing 11 families, 26,640 bacterial ASVs representing 332 families, and 6,306 fungal ASVs representing 240 families were detected (Fujita et al., 2024) (Figures 2; Figure 2–figure supplement 1).

**Figure 2.**
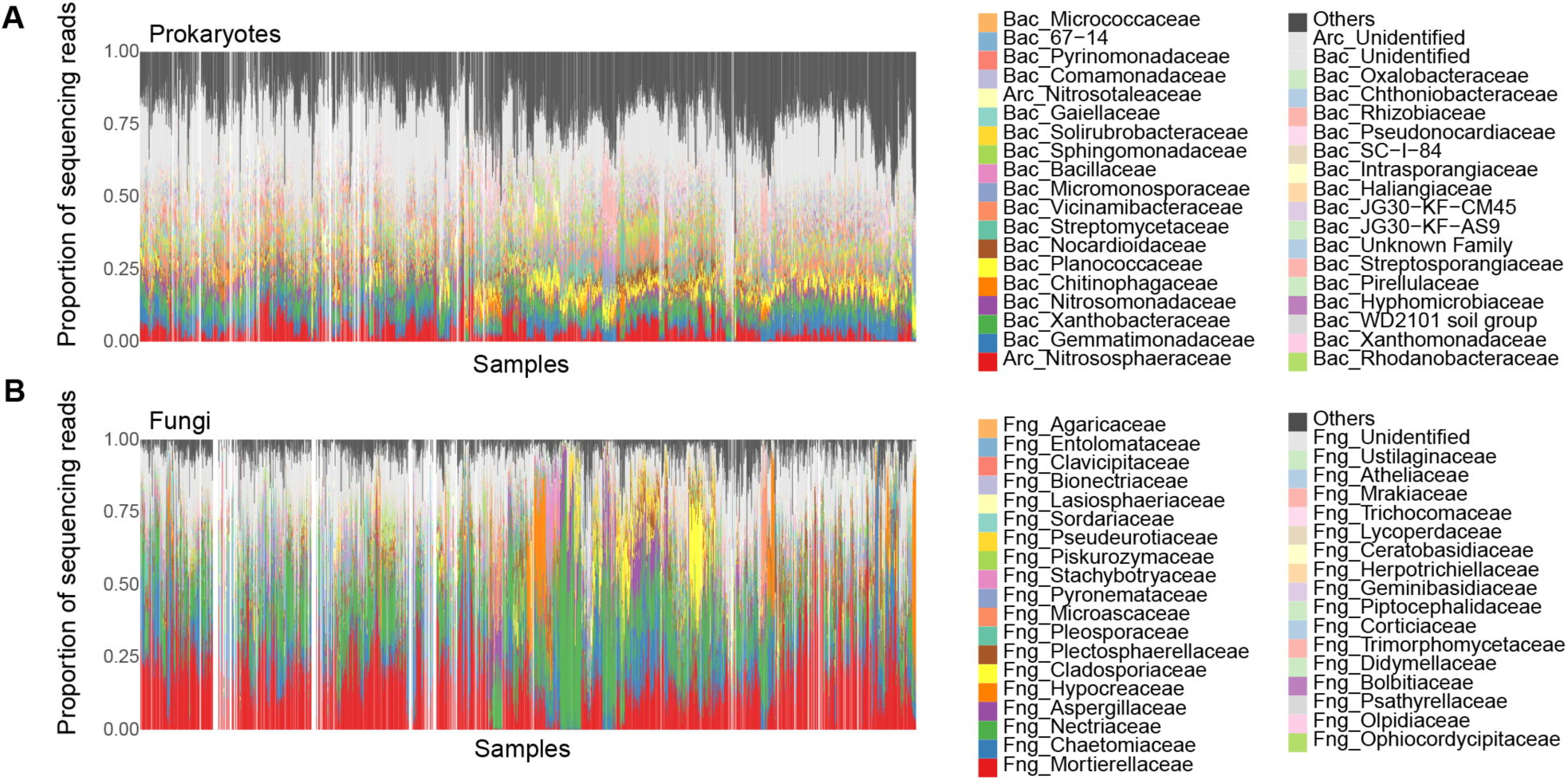
Community structure of the source data. The family-level compositions of prokaryotes (**A**) and fungi (**B**) are shown based on the source dataset (Fujita et al., 2024). The soil samples from which DNA sequence data were unavailable for either prokaryotic 16S rRNA or fungal ITS regions are indicated as blanks. **Figure supplement 1.** Community structure of the source data (order- and genus-level compositions).

### Community structure along environmental gradients

We first inspected how prokaryotic and fungal community structure varied along environmental gradients. For each data matrix representing the family-level compositions of prokaryotes or fungi, a principal coordinate analysis (PCoA) was performed based on Bray-Curtis *β*-diversity. The PCoA1 and PCoA2 scores were then plotted, respectively, along the axes of soil environmental factors. Specifically, the axes of the environmental factors were defined based on a principal component analysis (PCA) of soil pH, electrical conductivity, C/N ratio, and available phosphorous concentration. In total, 1,771 and 1,664 samples for which the information of both community structure and all the four environmental variables was available were included in the analyses of prokaryotes and fungi, respectively. For each plot representing relationship between environmental conditions and community structure, the density of data points was visualized with the ggplot2 3.3.6 package (Wickham, 2011) of R v.4.1.2 (R Core Team, 2020).

### Energy landscape analysis

We examined the stability landscape of soil microbiome structure based on the framework of an energy landscape analysis (H Fujita et al., 2023; Suzuki et al., 2021; Watanabe et al., 2014) (tutorials of energy landscape analyses are available at https://github.com/kecosz/rELA). In the framework, the term “energy” is defined by the following equations based on statistical physics (Suzuki et al., 2021; Watanabe et al., 2014). Within the “assembly graphs” representing paths of community dynamics (Coyte et al., 2021; Serván and Allesina, 2021), probabilities of observing specific community compositions can be explored as detailed previously (Suzuki et al., 2021). In brief, probabilities of community states 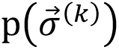 are given by

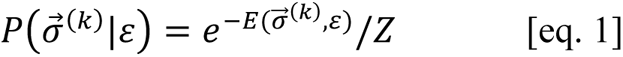

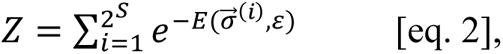

where 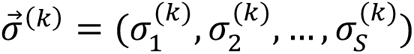 is a community state vector of *k*-th community state and *S* is the total number of taxa (e.g., ASVs, species, genera, or families) examined. *ε* = (*ε*_1_, *ε*_2_, ..., *ε_M_*) is an array of continuous values representing environmental factors (e.g., soil pH and electrical conductivity) and *M* is the total number of environmental parameters. *σ_i_*^(*k*)^ is a binary variable that represents presence (1) or absence (0) of taxon *i*: i.e., there are a total of 2*^s^* community states. As the exploration of the 2*^s^* community states were computationally intensive, we coded community states at the family-level taxonomic compositions. Specifically, for each sample, families whose relative abundance exceeds a certain threshold value (threshold for binarization) were coded as 1, while the remaining minor families were coded as 0. Subsequently, families whose occurrence ratios (i.e., the proportions of samples in which target families were coded as 1) were less than a certain threshold (occurrence threshold) were excluded from the dataset.

Likewise, families that appeared in almost all samples (1 – occurrence threshold) were excluded. Note that without such thinning of input data, the dimensions of community states are too high to be explored even using supercomputers. Therefore, exclusion of the taxa that contribute little to the classification of community states (i.e., taxa appearing only in a small fraction of samples or those appearing in most samples) is inevitable in the energy landscape analysis. Through intensive preliminary computational runs with various combinations of binarization and occurrence thresholds, we found that the number of taxa (*S*) should be kept less than 65 as detailed in the next subsection.

When input community matrix is defined, the energy of the community state 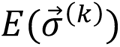 is given by the extended pairwise maximum entropy model:

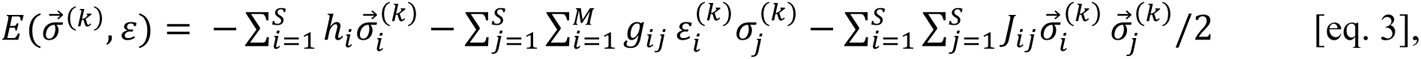

where *h_i_*, represents the net effect of implicit abiotic factors, by which *i*-th taxon is more likely to present (*hi*> 0) or not (*hi*< 0), *g_,ij_* represents the effect of the *i-*th observed environmental factor, and *j_,ij_* represents a co-occurrence pattern of *i*-th and *j*-th taxa. Since the logarithm of the probability of a community state is inversely proportional to 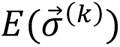, a community state having lower *E* is more frequently observed. To consider dynamics on an assembly graph defined as a network whose 2^s^ nodes represent possible community states and the edges represents transition path between them (two community states are adjacent only if they have the opposite presence/absence status for just one species), we assigned energy to nodes with the above equation, and so imposed the directionality in state transitions. Then, by using the steepest descent algorithm (Suzuki et al., 2021), we identified nodes having the lowest energy compared to all its neighbors within the weighted network, and determined their basins of attraction (Lewontin, 1969; Suzuki et al., 2021). These community states whose energy was lower than that of all adjacent community states represent estimated point equilibria (attractors), around which community states are expected to show transient fluctuations due to demographic stochasticity as considered in the statistical framework (H Fujita et al., 2023; Suzuki et al., 2021) (Figure 1). Soil pH, electrical conductivity, C/N ratio, and available phosphorous concentration were included as environmental variables in the model after normalization within the ranges from 0 to 1.

### Energy landscape structure

The energy landscapes of community structure were inferred, respectively, for three types of datasets, namely, the prokaryotic community matrix, the fungal matrix, and the matrix including both prokaryotes and fungi. As mentioned above, various combinations of binarization and occurrence thresholds were examined to check the reproducibility of the results. In addition to the energy landscape analysis based on the above-mentioned family-level delineation of community states, analyses based on community-state delineation at the order-level were performed. In the main body and supplementary figures of this study, we show the results at the following settings: prokaryotes (family), binarization = 0.020, occurrence = 0.10; prokaryotes (order), binarization = 0.020, occurrence = 0.10; fungi (family), binarization = 0.001, occurrence = 0.05; fungi (family), binarization = 0.001, occurrence = 0.10; prokaryotes + fungi (family), binarization = 0.030, occurrence = 0.10; prokaryotes + fungi (order), binarization = 0.030, occurrence = 0.10. Note that these thresholds were selected to make the state space (2*^s^*) neither too simplified (e.g., *S* < 30) nor too complex (*S* < 65).

For each setting, the parameters of the extended pairwise maximum entropy model [eq. 3] were adjusted to the empirical data. More precisely, the maximum likelihood estimates of *h_i_*, *g_ij_*, and *j_ij_* was obtained by a stochastic approximation method as detailed elsewhere (Suzuki et al., 2021). The parameters were regularized by a logistic prior with location 0 and scale 2.0 (for environmental responses) or 0.5 (for pairwise relationships) (Harris, 2016). Hyperparameters for the algorithm, criterion value for judging the convergence of parameters qth = 10^-5^, were set according to a series of preliminary analyses. Based on the inferred maximum entropy model, we determined basins of attraction (Lewontin, 1969) within the energy landscape based on a steepest descent procedure (Suzuki et al., 2021). The structure of the energy landscape was visualized by showing the energy of each soil sample on the two-dimensional surface of the community state space defined with the abovementioned PCoA scores. The default setting of environmental variables (the mean value for each of soil pH, electrical conductivity, C/N ratio, and available phosphorous concentration) was used in the energy calculation. Spline smoothing of the energy landscape was performed with optimized penalty scores using the mgcv v.1.8-40 package (Wood, 2022) of R. For each analysis of the prokaryote, fungi, and prokaryote + fungi datasets, 1,771, 1,664, and 1,474 samples for which the information of both community structure and all the four environmental variables was available were subjected to the analysis, respectively.

### Ecosystem functions and key taxa

For the inferred basins of microbial community compositions, associations with crop disease prevalence were examined. We first constructed the list of soil samples whose community structure was located within each basin of attraction. We then evaluated the ecosystem-scale properties of the basins in light of the metadata of crop disease symptoms (Fujita et al., 2024). Specifically, for each basin, we calculated the proportion of constituent soil samples with the minimal level of crop disease symptoms (the percentage of diseased plants < 20 or disease severity index < 20; (Fujita et al., 2024)). The bottoms of basins representing different levels of crop disease prevalence were then compared in terms of taxonomic compositions in order to explore microbial taxa that were keys to distinguish potentially disease-suppressive and disease-promotive soil ecosystems.

### Disconnectivity graphs

For the reconstructed energy landscape, we inferred “disconnectivity graphs” (Suzuki et al., 2021) representing how basins of attraction were split by tipping points (Figure 1A). Within a disconnectivity graph, community states whose energy is much lower than the energy of connected tipping points are expected to be resistant to perturbations (demographic stochasticity). In contrast, community states with small energy gaps to tipping points may be shifted from current basins to adjacent basins with minimal perturbations.

## Results

### Community structure along environmental gradients

On each plot showing community compositions (PCoA1 or PCoA2 scores) along the soil environmental gradient (Figure 3), multiple clusters of data points were observed for both prokaryotes and fungi (Figure 3–figure supplements 1-2). In other words, community states are expected to be classified into some clusters even under equivalent edaphic conditions.

**Figure 3.**
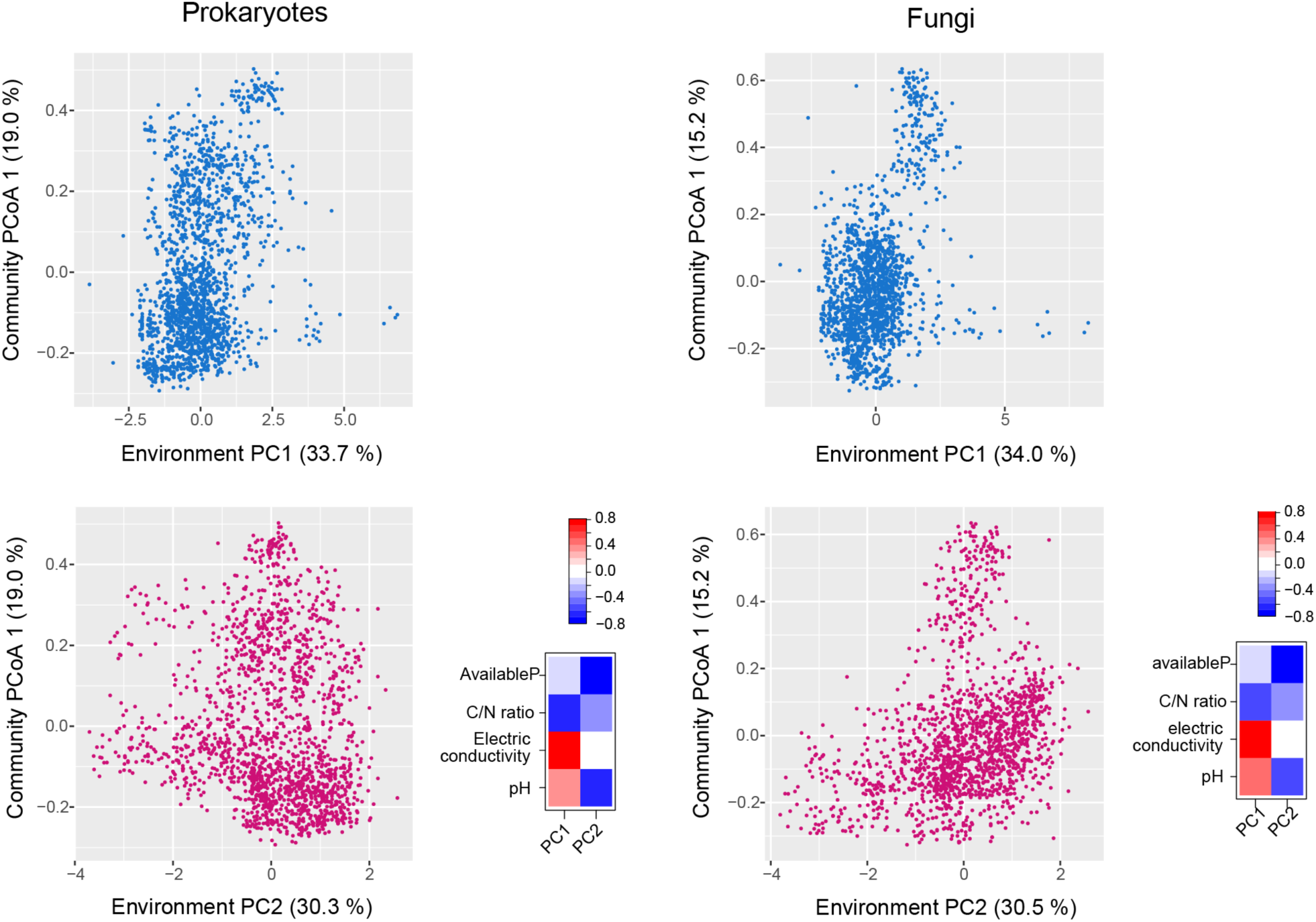
Community structure along environmental gradients. The scores representing prokaryotic/fungal community compositions (community PCoA1 scores) are shown along each PCA axis of soil environmental conditions. Regarding the environmental PCA axes, factor loadings of environmental variables examined (pH, electrical conductivity, C/N ratio, and available phosphorous concentration) are shown separately for prokaryotic (*N* = 1,771) and fungal (*N* = 1,664) datasets. **Figure supplement 1.** Prokaryotic community structure along environmental gradients (detailed analyses). **Figure supplement 2.** Fungal community structure along environmental gradients (detailed analyses).

### Energy landscape structure

The energy landscape of the family-level prokaryotic data included several major basins differing remarkably in associations with the prevalence of crop plant disease (Figure 4). Specifically, 59.6% of soil samples located within a basin (basin ID = 0IK1G2) were associated with the minimal plant-disease level, while the proportion was only 10.7% for another basin (LQWZ02) (Figure 4C-D). The presence of basins differing greatly in their associations with plant-disease levels was inferred as well at the order-level analysis of the prokaryotic data (Figure 4–figure supplement 1). Such variation in crop disease prevalence among inferred basins was observed also for the family-level analysis of fungal community structure (Figure 5). Specifically, while 57.9% of samples belonging to the basin 7QH9moTf8Xa, but none of the samples belonging to the basin 68C0849W020, were associated with the minimal plant-disease level (Figure 5D). Meanwhile, such difference in associations with disease prevalence was moderate in an analysis in which a smaller number of fungal families were examined to define community states (Figure 5–figure supplement 1). The presence of multiple basins, which differed in associations with crop-disease prevalence, was suggested even when prokaryotic and fungal community data were simultaneously analyzed (Figure 4–figure supplements 2-3).

**Figure 4.**
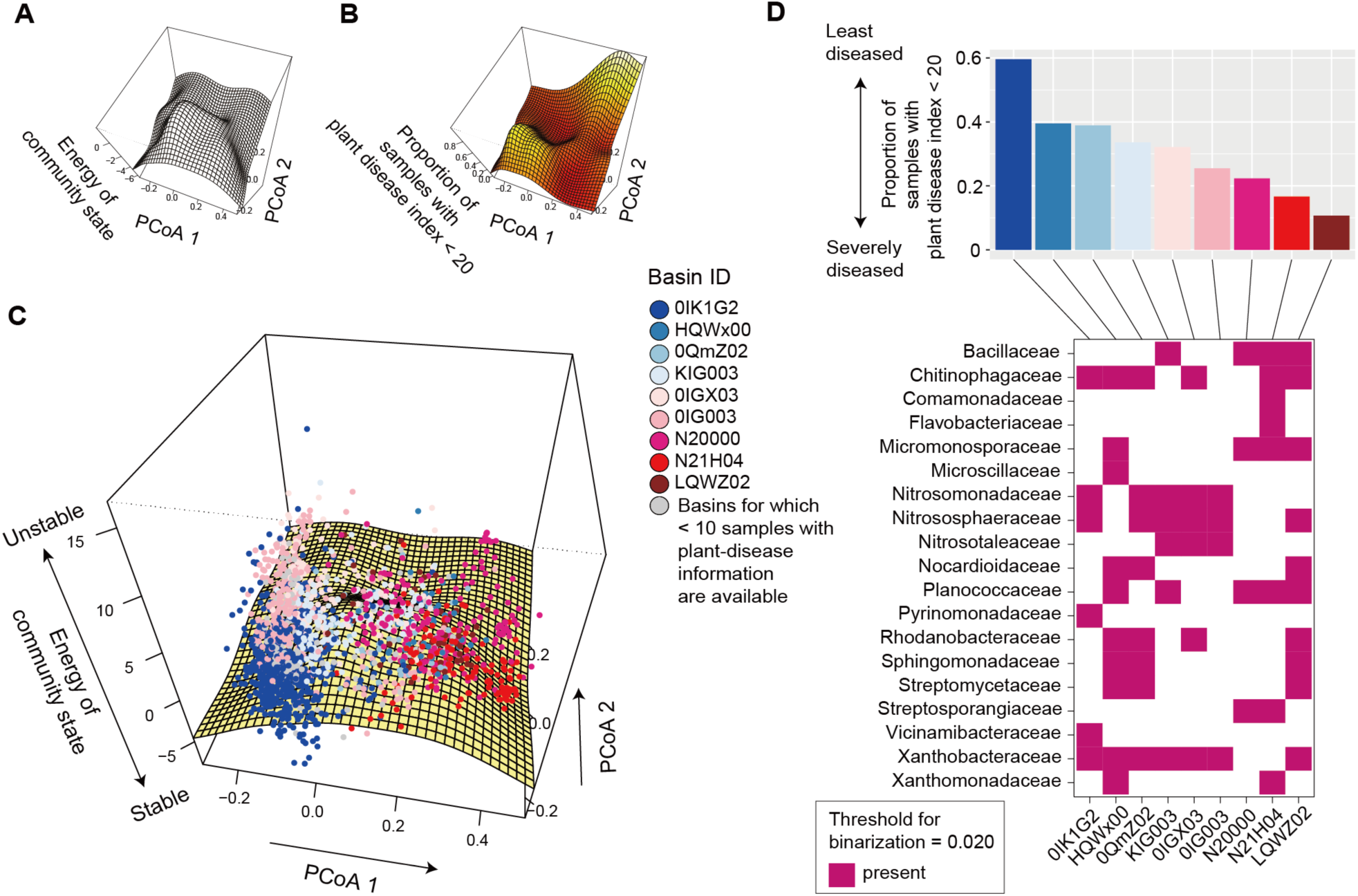
Energy landscape of prokaryotic communities. (**A**) Inferred energy landscape of family-level prokaryotic community structure (threshold for binarization = 0.020; occurrence threshold = 0.10; *S* = 35). The surface of energy levels was reconstructed across the PCoA space of fungal community structure (community PCoA1 and PCoA2 scores in Figures 2–figure supplement 1) based on spline smoothing. Community states with lower energy are inferred to be more stable. (**B**) Landscape of crop disease prevalence. Across the PCoA space of prokaryotic compositions, the proportion of samples with disease severity index < 20 is shown based on spline smoothing. (**C**) Community data points on the energy landscape. The axis of “energy of community state” is more expanded than that in panel **A** in order to cover the range of samples. Data points (samples) indicated by the same color belong to the same basins of attraction, which are represented by the IDs of the community states whose energy is lower than that of any adjacent community states (i.e., bottoms of basins). (**D**) Key taxa whose abundance represent basins. In the upper panel, the mean proportion of soil samples with the minimum level of plant (crop) disease symptoms (the percentage of diseased plants < 20 or disease severity index < 20) is shown for each basin. The lower panel indicates the key taxa whose abundance characterizes difference among the bottoms of the basins. **Figure supplement 1.** Energy landscape of prokaryotic communities (order-level compositions; threshold for binarization = 0.020; occurrence threshold = 0.10; *S* = 32). **Figure supplement 2.** Energy landscape of communities including both prokaryotes and fungi (family-level compositions; threshold for binarization = 0.030; occurrence threshold = 0.10; *S* = 31). **Figure supplement 3.** Energy landscape of communities including both prokaryotes and fungi (order-level compositions; threshold for binarization = 0.030; occurrence threshold = 0.10; *S* = 32).

**Figure 5.**
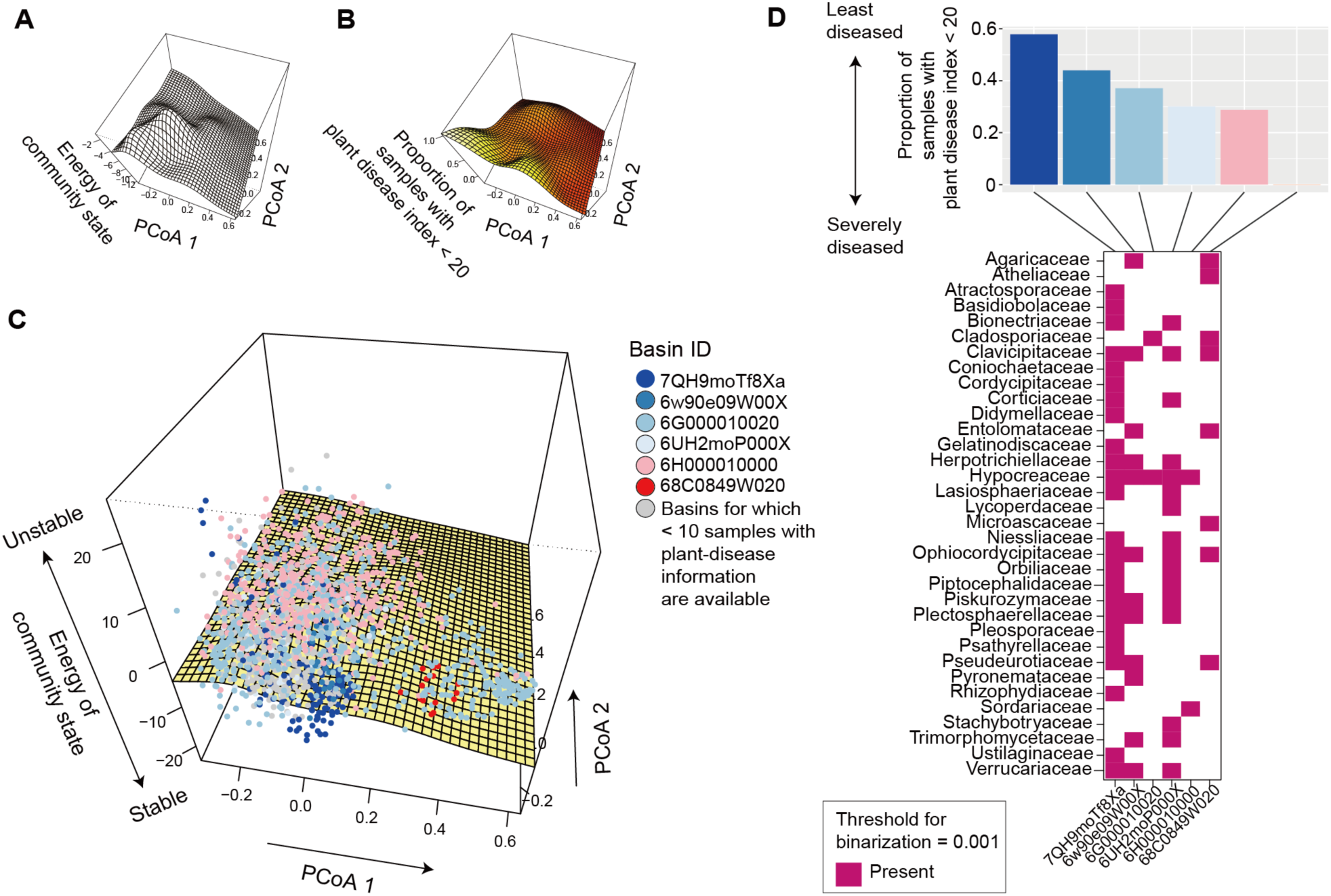
Energy landscape of fungal communities. (**A**) Inferred energy landscape of family-level fungal community structure (threshold for binarization = 0.001; occurrence threshold = 0.05; *S* = 62). The surface of energy levels was reconstructed across the PCoA space of fungal community structure (community PCoA1 and PCoA2 scores in in Figures 2–figure supplement 2) based on spline smoothing. Community states with lower energy are inferred to be more stable. (**B**) Landscape of crop disease prevalence. Across the PCoA space of prokaryotic compositions, the proportion of samples with disease severity index < 20 is shown based on spline smoothing. (**C**) Community data points on the energy landscape. The axis of “energy of community state” is more expanded than that in panel **A** in order to cover the range of samples. Data points (samples) indicated by the same color belong to the same basins of attraction, which are represented by the IDs of the community states whose energy is lower than that of any adjacent community states (i.e., bottoms of basins). (**D**) Key taxa whose abundance represent basins. In the upper panel, the mean proportion of soil samples with the minimum level of plant (crop) disease symptoms (the percentage of diseased plants < 20 or disease severity index < 20) is shown for each basin. The lower panel indicates the key taxa whose abundance characterizes difference among the bottoms of the basins. **Figure supplement 1.** Energy landscape of fungal communities (family-level compositions; threshold for binarization = 0.001; occurrence threshold = 0.10; *S* = 42).

### Ecosystem functions and key taxa

In an analysis of the prokaryotic community structure, 19 families were keys to distinguish basins differing in associations with crop-disease prevalence (Figure 4D). The presence of Pyrinomonadaceae and Vicinamibacteraceae, for example, was unique to the basin with the highest proportion of samples showing the minimal plant-disease level (Figure 4D). Likewise, in an analysis of the fungal community structure, the basin associated closely with the minimal plant-disease prevalence (7QH9moTf8Xa) was defined by the presence of several families such as Basidiobolaceae, Cordycipitaceae, and Gelatinodiscaceae (Figure 5D). The exploration of microbial taxa keys to distinguish basins with different ecosystem-level functions can be performed at other taxonomic levels (e.g., order-level; Figures 4–figure supplements 1 and 3).

### Disconnectivity graphs

Within the energy landscape of the family-level analysis of prokaryotes (Figure 4), both the basins associated with the least-diseased (OIK1G2) and most-diseased (N21H04) crop status were the deepest among the inferred basins (i.e., showing the largest energy gaps from the bottom to tipping points; Figure 6A-B). In the family-level analysis of fungi, the basin associated with the least-diseased status (7QH9moTf8Xa) was the deepest, while the other basin representing the most-diseased status (68C0849W020) was the shallowest (Figure 6C).

**Figure 6.**
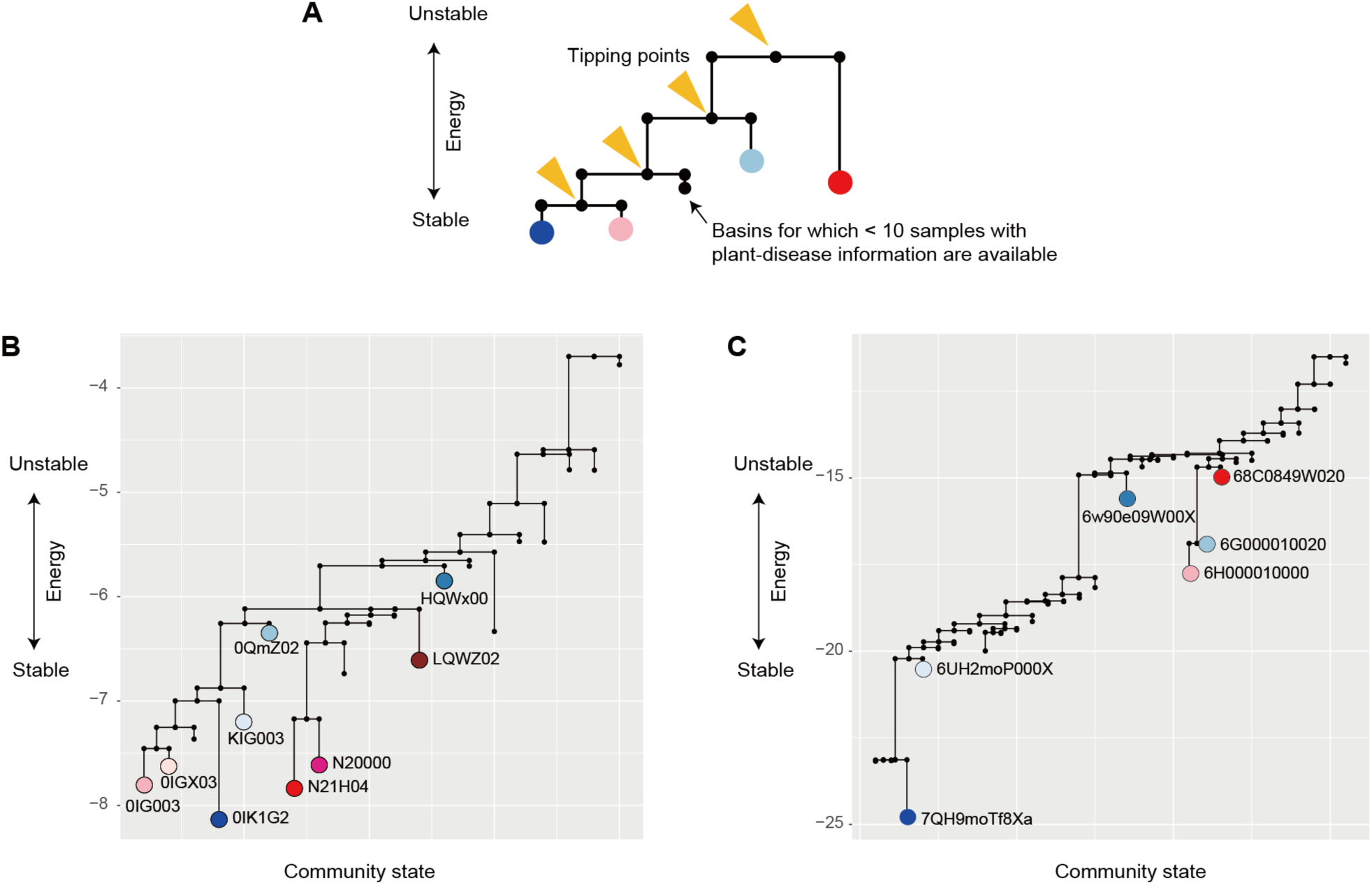
Disconnectivity graphs of the energy landscapes. (**A**) Schema of a disconnectivity graph. The energy of the “tipping points” splitting basins of attraction are presented across the axis of 2*^S^* possible community states, where *S* denotes the number of the species or taxa examined. The energy of the bottom of each basin is shown. (**B**) Tipping points and basins on the energy landscape of prokaryotes. The major basins of attraction with ζ 10 samples with plant-disease information are highlighted with the colors defined in Figure 4. (**C**) Tipping points and basins on the energy landscape of fungi. The major basins of attraction with ζ 10 samples with plant-disease information are highlighted with the colors defined in Figure 5.

## Discussion

We have estimated the stability landscape structure of complex microbiomes based on a statistical framework commonly applicable to diverse types of biological communities. The energy landscape analysis allows systematic analyses of taxon-rich community datasets by incorporating the information of multiple environmental factors (Dakos and Kéfi, 2022; Sánchez-Pinillos et al., 2024; Suzuki et al., 2021). While classic studies on community multistability have discussed ecological processes spanning a few intuitively distinguishable community states [high/low tree cover in forest-savanna transitions (Hirota et al., 2011; Staver et al., 2011a, 2011b) or macrophyte-/phytoplankton-dominated state in shallow lakes (Ibelings et al., 2007; Scheffer and Carpenter, 2003)], it is now made possible to define basins of attraction based on high-dimensional community datasets involving hundreds of species/taxa (Arumugam et al., 2011; Costea et al., 2017; H Fujita et al., 2023; Guim Aguadé-Gorgorió et al., 2023; Hayashi et al., 2024). Application of the general statistical platform will enhance our understanding of how stability landscape properties differ among diverse microbial and non-microbial systems.

Despite numerous potential compositions (2*^s^* community states; *S* is the number of considered species/taxa), the prokaryotic and fungal community states were grouped into small numbers of basins within energy landscapes (Figures 4-5). This result suggests that soil microbiome structure remain within certain regions even after demographic perturbations. In other words, once trapped in a basin of attraction, large shifts in community structure would not occur without perturbations whose strength exceed certain thresholds (Beisner et al., 2003; Lewontin, 1969; May, 1977; Scheffer et al., 1993). Importantly, the threshold strength of perturbations is estimated as the energy gap between bottoms of basins and tipping points (Suzuki et al., 2021) (Figure 6A). Furthermore, potential paths of community structural transitions can be quantitatively inferred as illustrated in disconnectivity graphs (Suzuki et al., 2021) (Figures 6B-C). Such statistical framework of quantitative science will entail novel opportunities for testing theories on biological community processes in the era of massive datasets.

Among potential processes or mechanisms underlying the multistability of community structure, historical contingency is of particular interest (Fukami, 2015). In the local assembly of microbial communities, early colonizers or residents can prevent the settlement of followers by constructing physical barriers (e.g., biofilms and mycelia) (Baümler and Sperandio, 2016; Fukami, 2015; Leopold et al., 2017; Verbruggen et al., 2013; Werner and Kiers, 2015) or emitting antibiotics (Mendes et al., 2013; Raaijmakers et al., 2002). In addition to those antagonistic effects on late colonizers, webs of mutualistic or commensalistic interactions within the microbiomes of early colonizers (Elias and Banin, 2012; Hiroaki Fujita et al., 2023; Zelezniak et al., 2015) would influence community dynamics. Due to such “priority effects” (Fukami, 2015), bacterial and fungal community compositions may persist within limited ranges of community states without substantial perturbations. Given that abilities to form physical or chemical barriers can differ greatly among microbial species/taxa (Mendes et al., 2013; Raaijmakers et al., 2002; Werner and Kiers, 2015), such variation in constituent species’ priority effects may underly the observed variation in the depth of basins (Figure 6B-C).

The inference of stability landscape structure provided an opportunity for evaluating relationship between community stability and ecosystem-scale functions. The basins of attraction of prokaryotic/fungal community structure differed considerably in associations with crop disease prevalence (Figure 5), suggesting the presence of “stable and favorable” and “stable but unfavorable” states of microbiomes (Mendes et al., 2011; Schlatter et al., 2017; Yuan et al., 2020) in terms of agricultural productivity. This finding adds an important dimension of discussion on the use of microbes in agriculture. Beyond investigations on single species/strains of microbes, microbiome studies have explored sets of microbes that collectively maximize biological functions (Jansson and Hofmockel, 2018; Toju et al., 2018; Trivedi et al., 2020; Vorholt et al., 2017). In particular, experimental studies on “synthetic” communities have reorganized our knowledge of microbiome functions (Jansson and Hofmockel, 2018; Trivedi et al., 2020; Vorholt et al., 2017). Nonetheless, such microbial functions cannot be realized in real agroecosystems if the synthesized or designed microbiome compositions are vulnerable to biotic and abiotic environmental changes in the wild (Mazzola and Freilich, 2017). Thus, in addition to functional properties, compositional stability is the key to manage microbiomes in agroecosystems (Faust and Raes, 2012; Toju et al., 2020; Vorholt et al., 2017).

In our analysis across the Japan Archipelago, prokaryotic and fungal taxa keys to distinguish least-diseased and severely-diseased states of soil microbiomes were highlighted (Figures 4-5). Among them, Basidiobolaceae and Cordycipitaceae are of particular interest because they include many species potentially utilized as biological control agents for suppressing pest insects (Meyling and Eilenberg, 2007; Möckel et al., 2022). Gelatinodiscaceae is another fungal taxon playing potentially important roles as symbionts of plants (Johnston et al., 2019). These results illuminate the hypothesis that plant disease could be suppressed under the coexistence of multiple prokaryotic and fungal taxa with favorable ecosystem functions (Toju et al., 2018; Toju and Tanaka, 2019). Thus, statistical analyses of stability landscapes allow the exploration of key species or taxa (Paine, 1966; Power et al., 1996), whose management could result in transitions from unfavorable ecosystem states to favorable ones (Gunderson, 2000; Scheffer et al., 2001, 1993; Scheffer and Carpenter, 2003). Given that most prokaryotic and fungal families highlighted in our analysis have cosmopolitan distributions, a next crucial step is to test whether the basins defined across the Japan Archipelago can be used to categorize disease-suppressive and disease-susceptible microbiomes in other regions on the globe.

Although the energy landscape analysis enhances our understanding of community stability and functions, its results should be interpreted with caution. First, given that classic empirical studies examined community multistability with system-specific simple criteria [e.g., high/low tree cover (Hirota et al., 2011; Staver et al., 2011a, 2011b)], special care should be taken when we extend the approach to species-rich (high-dimensional) community datasets (Guim Aguadé-Gorgorió et al., 2023). In other words, unambiguous and broadly applicable criteria based on statistical evaluation are the prerequisite for comparative analyses of community multistability.

Although we applied a straightforward statistical definition of basins of attraction (Suzuki et al., 2021) (Figure 1) in light of classic theoretical studies (Beisner et al., 2003; Lewontin, 1969; May, 1977; Scheffer et al., 1993), continuous methodological improvements should be explored towards further comprehensive analyses. Second, our analysis on hyper-diverse soil microbiomes incurred substantial computational costs, forcing us to limit the energy landscape analysis to family-level input data. Further improvements of codes are necessary for inferring stability landscapes at genus-, species-, or strain-level analyses. Third, it should be acknowledged that detailed discussion on ecological processes require time-series datasets (Davidson et al., 2023; Scheffer et al., 2012, 2009). Because our present data lacked the information of temporal changes in community structure, we are unable to discuss the frequency and pace of community structural transitions between basins of attraction. Monitoring of microbiome compositions (Faust et al., 2015; Hayashi et al., 2024; Yajima et al., 2023) is necessary for filling the gap between theoretical and empirical studies (Long et al., 2024).

The energy landscape framework of multistability analysis is readily applicable to a wide range of microbiome datasets. Application to human microbiome data is of particular interest in terms of the confirmation of the existence of multiple basins of attraction (Jeffery et al., 2012). In addition, insights into the key microbial species/taxa that would play key roles in the transitions from disease-associated microbiome states to healthy ones will open new directions of microbiome therapy. Furthermore, time-series analyses of community dynamics on stability landscapes will allow us to forecast and prevent transitions into unfavorable community states [e.g., dysbiosis (Carding et al., 2015; H Fujita et al., 2023; Long et al., 2024)]. Along with such extensions of observational research, experimental studies controlling key species/taxa or environmental parameters (Schröder et al., 2005) will promote both basic and applied sciences of ecosystem functions.

## Acknowledgements

We thank the SuperComputer System, Institute for Chemical Research, Kyoto University for the use of super computers.

## Author contributions

H.F., S.Y. and H.T. designed the work. H.F. performed molecular experiments. H.F. and H.T. analyzed the data. H.T. wrote the paper with H.F., S.Y., and K.S.

## Data accessibility

The accession number of the DDBJ Sequence Read Archive: DRA015491. The microbial community matrices are provided with codes at our GitHub repository (https://github.com/hiro-toju/Soil_EnergyLandscape_NARO3000) [to be released after the acceptance of the manuscript].

## Notes

**Funding** This work was financially supported by JST PRESTO (JPMJPR16Q6), JST FOREST (JPMJFR2048), Human Frontier Science Program (RGP0029/2019), JSPS Grant-in-Aid for Scientific Research (20K20586), NEDO Moonshot Research and Development Program (JPNP18016), and JST CREST (JPMJCR23N5) to H.T., JSPS Grant-in-Aid for Scientific Research (20K06820 and 20H03010) to K.S., and JSPS Fellowship to H.F.

### Summary of Updates

The terminology within the manuscript has been improved.

