## Supplementary Figures for "Basins of attraction of microbiome structure and soil ecosystem functions"

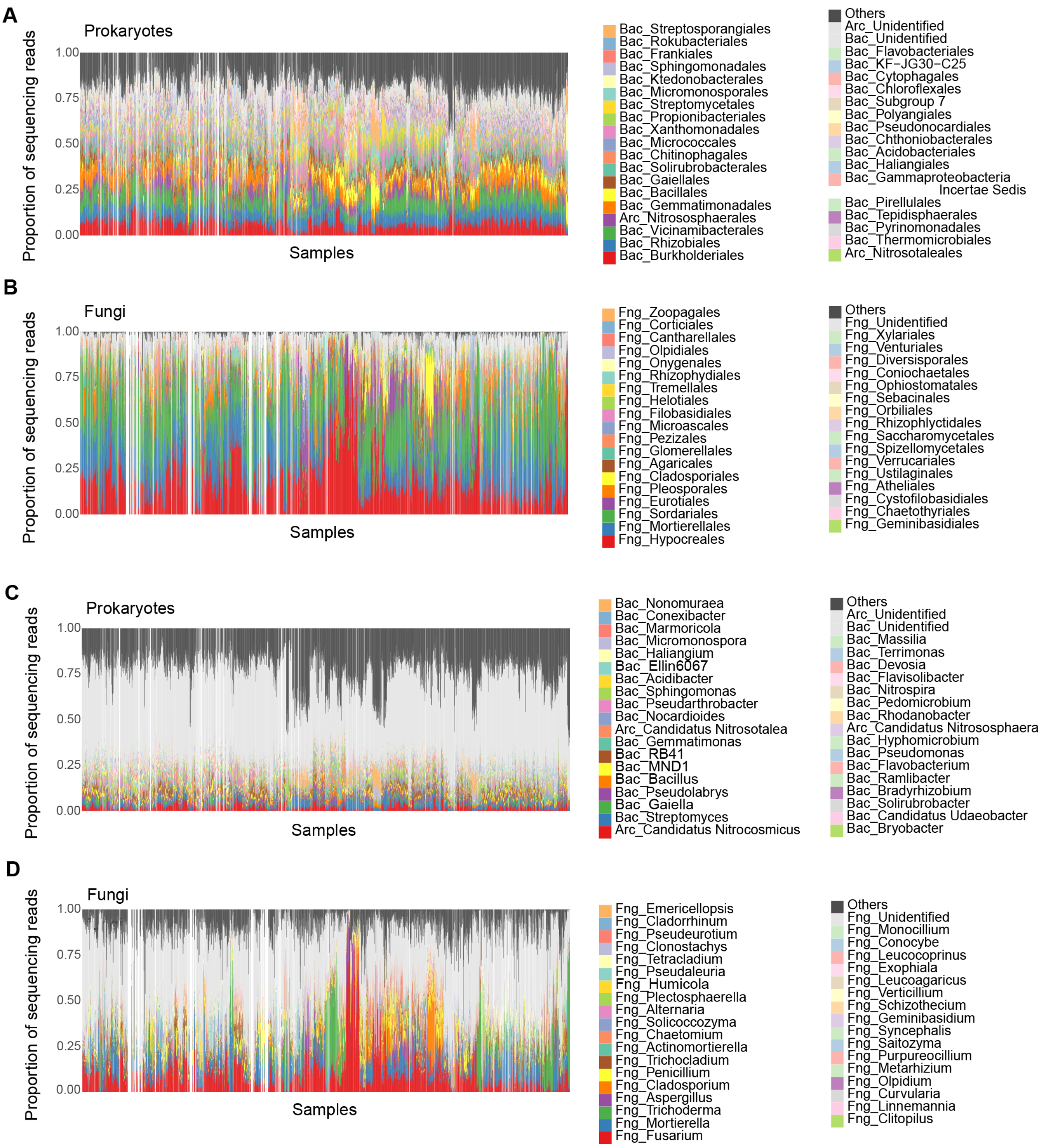


**Figure 2–supplement 1.** Community structure of the source data (order- and genus-level compositions). (**A**) Order-level compositions of prokaryotic communities. (**B**) Order-level compositions of fungal communities. (**C**) Genus-level compositions of prokaryotic communities. (**D**) Genus-level compositions of fungal communities. The soil samples from which DNA sequence data were unavailable for either prokaryotic 16S rRNA or fungal ITS regions are indicated as blanks.

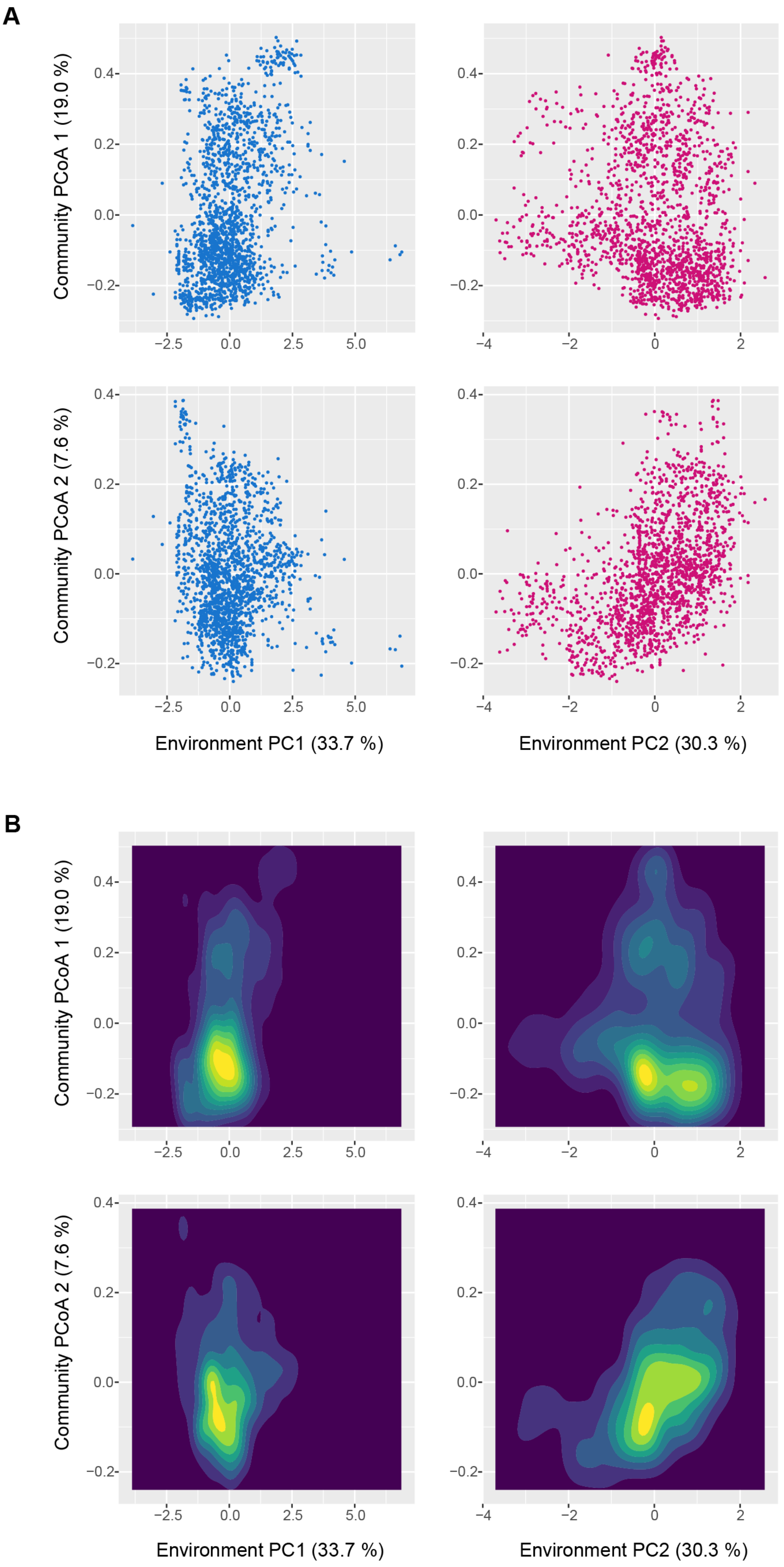


**Figure 3–supplement 1.** Prokaryotic community structure along environmental gradients (detailed analyses). The scores representing prokaryotic/fungal community compositions

(community PCoA 1 and 2 scores) are shown along each PCA axis of soil environmental factors. Regarding the environmental PCA axes, factor loadings of environmental variables examined (pH, electrical conductivity, C/N ratio, and available phosphorous concentration) are shown in Figure 3. (**A**) Scatter plots. (**B**) Density plots.

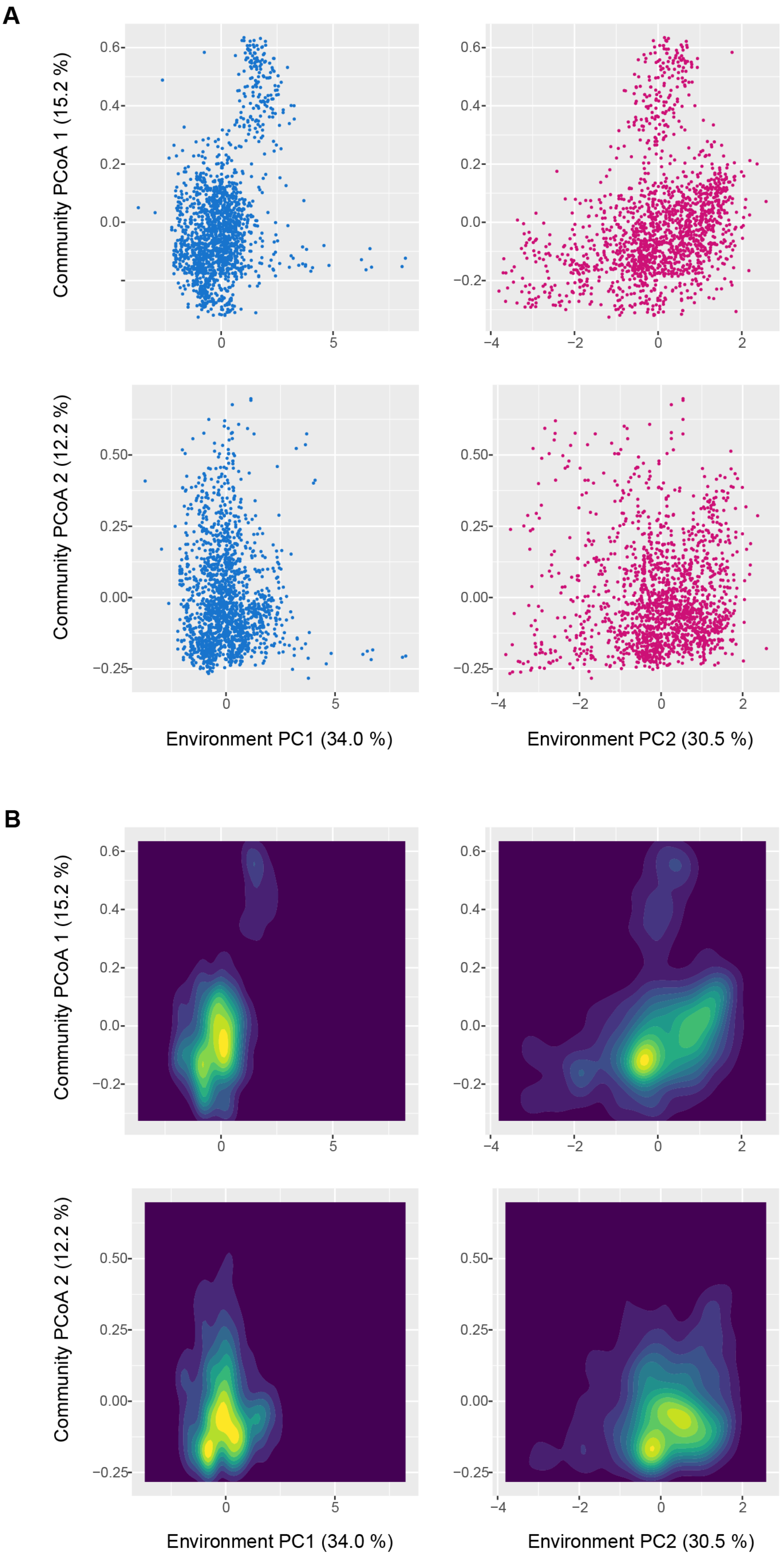


**Figure 3–supplement 2.** Fungal community structure along environmental gradients (detailed

analyses). The scores representing prokaryotic/fungal community compositions (community PCoA 1 and 2 scores) are shown along each PCA axis of soil environmental conditions. Regarding the environmental PCA axes, factor loadings of environmental variables examined (pH, electrical conductivity, C/N ratio, and available phosphorous concentration) are shown in Figure 3. (**A**) Scatter plots. (**B**) Density plots.

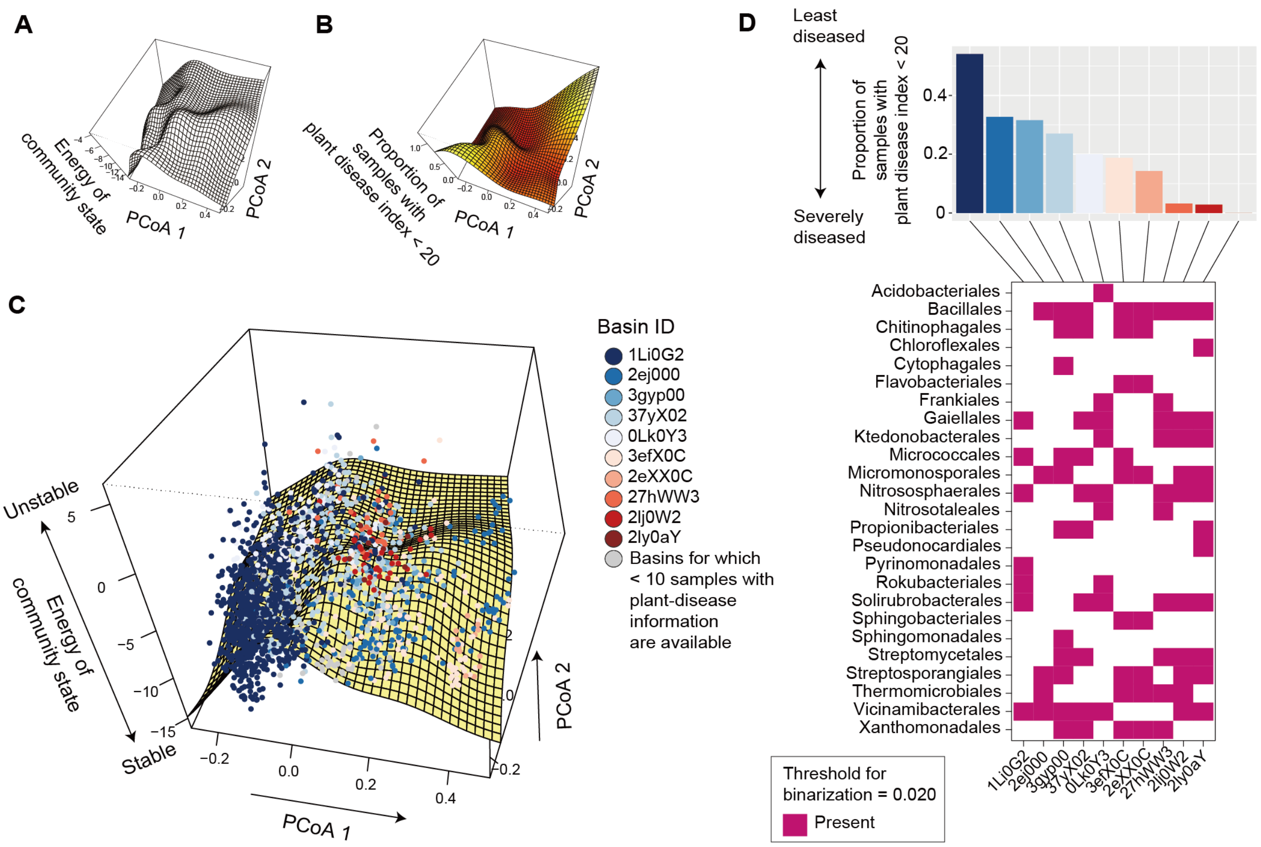


**Figure 4–supplement 1.** Energy landscape of prokaryotic communities (order-level compositions; threshold for binarization = 0.020; occurrence threshold = 0.10; $S = 32$). (**A**) Inferred energy landscape of order-level prokaryotic community structure. The surface of energy levels was reconstructed across the PCoA space of fungal community structure (community PCoA1 and PCoA2 scores in in Figures 2–figure supplement 1) based on spline smoothing. Community states with lower energy are inferred to be more stable. (**B**) Landscape of crop disease prevalence. Across the PCoA space of prokaryotic compositions, the proportion of samples with disease severity index < 20 is shown based on spline smoothing. (**C**) Community data points on the energy landscape. The axis of "energy of community state" is more expanded than that in panel **A** in order to cover the range of samples. Data points (samples) indicated by the same color belong to the same basins of attraction, which are represented by the IDs of the community states whose energy is lower than that of any adjacent community states (i.e., bottoms of basins). (**D**) Key taxa whose abundance represent basins. In the upper panel, the mean proportion of soil samples with the minimum level of plant (crop) disease symptoms (the percentage of diseased plants < 20 or disease severity index < 20) is shown for each basin. The lower panel indicates the key taxa whose abundance characterizes difference among the bottoms of the basins.

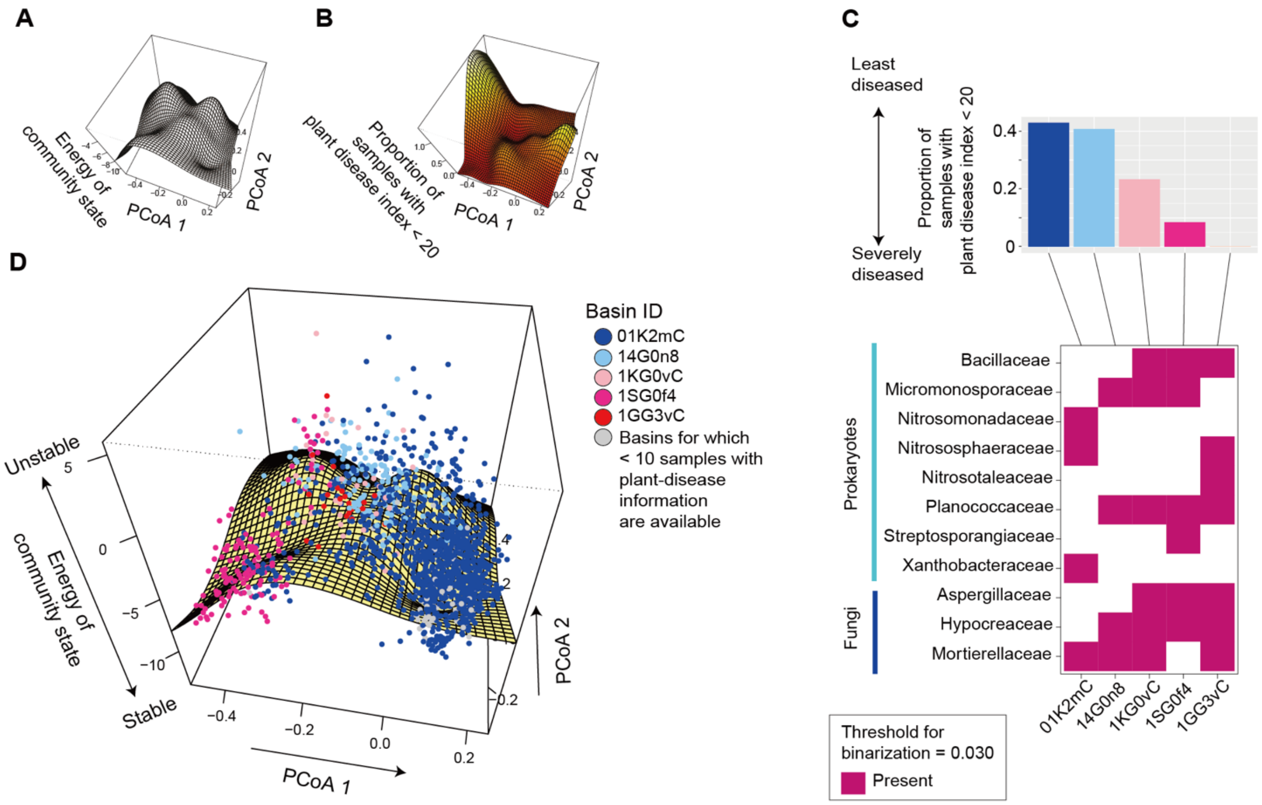


**Figure 4–supplement 2.** Energy landscape of communities including both prokaryotes and fungi (family-level compositions; threshold for binarization = 0.030; occurrence threshold = 0.10; $S$ = 31). (**A**) Inferred energy landscape. The surface of energy levels was reconstructed across the PCoA space of community structure (community PCoA1 and PCoA2 scores of the dataset including both prokaryotes and fungi) based on spline smoothing. Community states with lower energy are inferred to be more stable. (**B**) Landscape of crop disease prevalence. Across the PCoA space of prokaryotic compositions, the proportion of samples with disease severity index < 20 is shown based on spline smoothing. (**C**) Community data points on the energy landscape. The axis of “energy of community state” is more expanded than that in panel **A** in order to cover the range of samples. Data points (samples) indicated by the same color belong to the same basins of attraction, which are represented by the IDs of the community states whose energy is lower than that of any adjacent community states (i.e., bottoms of basins). (**D**) Key taxa whose abundance represent basins. In the upper panel, the mean proportion of soil samples with the minimum level of plant (crop) disease symptoms (the percentage of diseased plants < 20 or disease severity index < 20) is shown for each basin. The lower panel indicates the key taxa whose abundance characterizes difference among the bottoms of the basins.

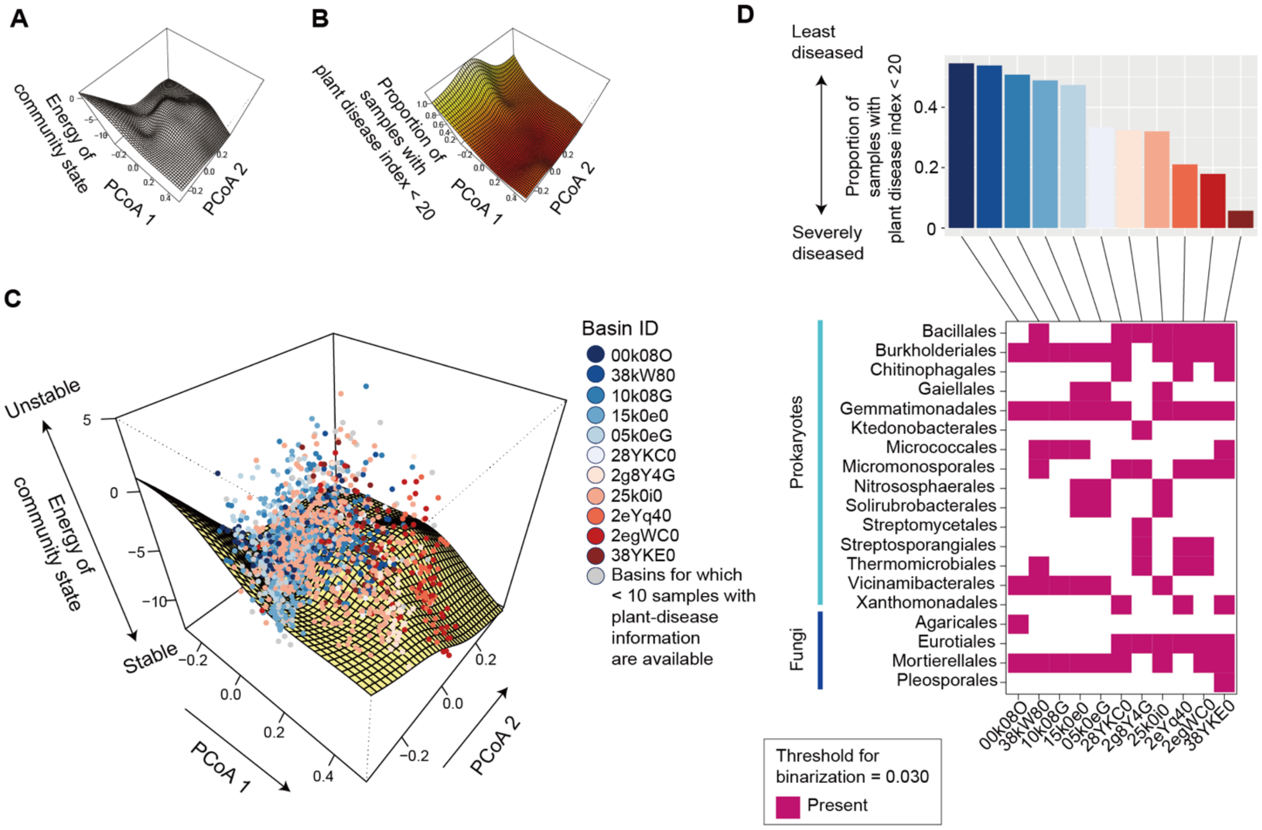


**Figure 4–supplement 3.** Energy landscape of communities including both prokaryotes and fungi (order-level compositions; threshold for binarization = 0.030; occurrence threshold = 0.10; $S$ = 32). (**A**) Inferred energy landscape. The surface of energy levels was reconstructed across the PCoA space of community structure (community PCoA1 and PCoA2 scores of the dataset including both prokaryotes and fungi) based on spline smoothing. Community states with lower energy are inferred to be more stable. (**B**) Landscape of crop disease prevalence. Across the PCoA space of prokaryotic compositions, the proportion of samples with disease severity index < 20 is shown based on spline smoothing. (**C**) Community data points on the energy landscape. The axis of "energy of community state" is more expanded than that in panel **A** in order to cover the range of samples. Data points (samples) indicated by the same color belong to the same basins of attraction, which are represented by the IDs of the community states whose energy is lower than that of any adjacent community states (i.e., bottoms of basins). (**D**) Key taxa whose abundance represent basins. In the upper panel, the mean proportion of soil samples with the minimum level of plant (crop) disease symptoms (the percentage of diseased plants < 20 or disease severity index < 20) is shown for each basin. The lower panel indicates the key taxa whose abundance characterizes difference among the bottoms of the basins.

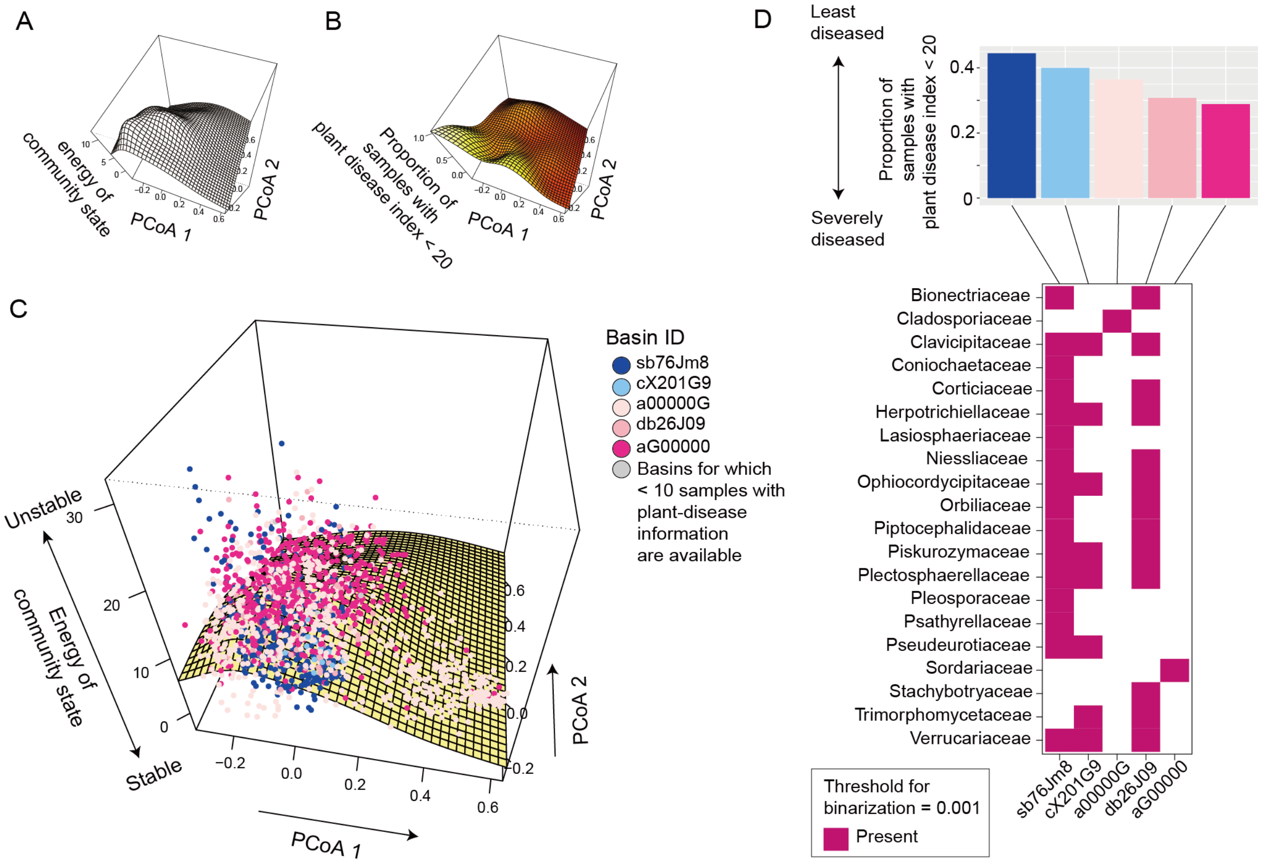


**Figure 5–supplement 1.** Energy landscape of fungal communities (family-level compositions; threshold for binarization = 0.001; occurrence threshold = 0.10; $S$ = 42). (**A**) Inferred energy landscape of family-level fungal community structure. The surface of energy levels was reconstructed across the PCoA space of fungal community structure (community PCoA1 and PCoA2 scores in in Figures 2–figure supplement 2) based on spline smoothing. Community states with lower energy are inferred to be more stable. (**B**) Landscape of crop disease prevalence. Across the PCoA space of prokaryotic compositions, the proportion of samples with disease severity index < 20 is shown based on spline smoothing. **(C)** Community data points on the energy landscape. The axis of “energy of community state” is more expanded than that in panel **A** in order to cover the range of samples. Data points (samples) indicated by the same color belong to the same basins of attraction, which are represented by the IDs of the community states whose energy is lower than that of any adjacent community states (i.e., bottoms of basins). (**D**) Key taxa whose abundance represent basins. In the upper panel, the mean proportion of soil samples with the minimum level of plant (crop) disease symptoms (the percentage of diseased plants < 20 or disease severity index < 20) is shown for each basin. The lower panel indicates the key taxa whose abundance characterizes difference among the bottoms of the basins.
